# Ischemic stroke and dietary vitamin B12 deficiency in old-aged females impaired motor function, increased ischemic damage size, and changed metabolite profiles in brain and cecum tissue

**DOI:** 10.1101/2022.04.04.487028

**Authors:** Joshua Poole, Paniz Jasbi, Agnes S. Pascual, Sean North, Neha Kwatra, Volkmar Weissig, Haiwei Gu, Teodoro Bottiglieri, Nafisa M. Jadavji

## Abstract

The global population is aging and the prevalence of age-related diseases, such as stroke, is predicted to increase. A vitamin B12 deficiency (vit. B12 def.) is common in the elderly, because of changes in metabolism. Clinical studies have reported that a vit. B12 def results in worse outcome after stroke, the mechanisms through which a vit. B12 def. changes the brain requires further investigation. This study investigated the role of vit. B12 def. on stroke outcome and mechanisms using aged female mice. Eighteen month old females were put on a control or vit. B12 def. diet for four weeks, after which an ischemic stroke was induced in the sensorimotor cortex. After damage, motor function was measured and animals were euthanized and tissues were collected for analysis. Vit. B12 def. animals had increased levels of total homocysteine in plasma and liver, choline levels were also increased in the liver. Vit. B12 def. animals had larger damage volume in brain tissue and more apoptosis. In the cecum, changes in creatinine and methylmalonic acid were observed in vit. B12 def. animals, pathway analysis showed dysfunction in B12 transport. Analysis of mitochondrial metabolomics in brain tissue showed reduced levels of metabolites involved in the TCA cycle in vit. B12 def animals. Meanwhile, pathway analysis showed significant, widespread dysfunction in phenylalanine, tyrosine, and tryptophan biosynthesis. Motor function after stroke was impaired in vit. B12 def. animals. A dietary vitamin B12 deficiency impairs motor function through increased apoptosis and changes in mitochondrial metabolism in brain tissue.

## Introduction

Stroke is among the leading causes of death globally and its prevalence as a major health concern is predicted to increase, as the global population ages and demographics of populations change (Mozaffarian et al., n.d.; Ovbiagele et al., 2013). Currently, stroke is prevalent and detrimental to elderly populations (> 65 years old) (Feigin et al., 2015; Ovbiagele et al., 2013). Ischemic stroke is the most common form of stroke. It is caused by blockage of arterioles leading to portions of the brain. The blockage results in reduced oxygen and energy supply to the brain causing severe disability and death. Many factors contribute to stroke risk and recovery outcomes, making it highly multifactorial. Nutrition is a modifiable risk factor for stroke (Mozaffarian et al., 2016.; Spence, 2019). For example, vitamin B12 deficiency is a well-established risk factor for stroke and poor stroke recovery outcomes. Approximately 20% of older adults (> 60 years old) have a vitamin B12 deficiency, making it of high concern to this population (Ahmed et al., 2019; Yahn et al., 2021).

^1^Cardiovascular disease (CVD) is the leading cause of death among men and women. Women have more risk factors and worse outcomes than men with CVD (Okunrintemi et al., 2018.). One of the many reasons these problems exist is that preclinical studies are targeted towards males (Persky et al., 2010). Over 90% of preclinical studies use strictly male mice whereas all clinical studies use equal part male and female participants (Persky et al., 2010; Cordonnier et al., 2017). This makes clinical pharmaceutical findings favor better outcomes in males, making female treatment worse than male (Macrae, 2011; Singh et al., 2021). This approach is taken in spite of the fact that stroke frequency and outcome in female mice and human participants changes depending age, menopause and other female-specific biological factors that are not applicable in males. Assessing these differences and strengthening female treatment is lost in the lack of female focus in preclinical studies (Persky et al., 2010; Singh et al., 2021). Another aim of this paper is to bridge this gap and increase insight into the female-specific stroke treatment.

Vitamin B12 is a component of one-carbon (1C) metabolism, which is a network that integrates nutritional signals with biosynthesis, redox homeostasis, and epigenetics and plays an essential role in the regulation of cell proliferation, stress resistance, and embryo development (Persky et al., 2010). A vitamin B12 deficiency results in increased levels of homocysteine, which is a risk for stroke. The literature has shown that patients with a vitamin B12 deficiency and hyperhomocysteinemia during an ischemic stroke have been reported to have worse outcome (Ahmed et al., 2019; Pieters et al., 2009; Zacharia et al., 2017). Vitamin B12 plays an essential role in mitochondrial energy production and cellular function (Depeint et al., 2006). Mitochondria are a major contributor to the development of apoptotic and necrotic cell death after ischemic stroke (Nguyen et al., 2018). The impact of vitamin B12 deficiency after stroke on mitochondrial function requires more investigation. The aim of this study was to investigate the impact of vitamin B12.

## Materials and methods

### Animals

All experiments in animals were approved by the Midwestern University IACUC committee. Female C57BL/6J mice were obtained from Jackson Laboratory for this study. Twenty-two 18-month-old female mice were obtained and, upon arrival, were habituated for 1 week prior to the start of experiments.

### Experimental Design

An overview of all experimental manipulations is outlined in Figure 7. After mice were habituated to the Midwestern University animal facility, mice were randomly assigned to control or vitamin B12 deficient groups and placed on diets for 4 weeks. At 18 months of age, ischemi stroke was induced in animals using the photothrombosis model (Abato et al., 2020; Jadavji et al., 2018, 2017; Labat-gest and Tomasi, 2013; Lee et al., 2004); this corresponds to middle-age in humans (Flurkey et al., 2007). The ischemic stroke damaged the sensorimotor cortex, which allows for motor function assessment. Four weeks after damage motor function was measured in animals using the forepaw placement, accelerating rotarod, and ladder beam tasks. After the completion of behavioral testing animals were euthanized and tissue was collected, including brain, liver, cecum, and blood tissue for further study.

### Diet

The mice were placed on a vitamin B12 deficient (vit. B12 def.) (0 mg/kg) or a control (0.025 mg/kg vitamin B12) diet four weeks prior to photothrombosis damage and four weeks post photothrombosis damage. The diets were formulated by and purchased from Envigo. The control diet (TD. 190790) contains the recommended dietary amount of nutrients for mice (Reeves, 1997). The vitamin B12 deficient diet (TD. 190791) was formulated based on a previous study in mice that has shown it to be safe and have no negative side effects (Bernard et al., 2018). The mice had *ad libitum* access to food and water throughout the experiment. Body weights of each animal were recorded weekly.

### Photothrombosis Model

Using the photothrombosis model of ischemic stroke damage, mice were anesthetized with 4-5% isoflurane in oxygen. After anesthetization, the mice had the top of their heads shaved and disinfected. Tear gel was used to prevent their eyes from drying out while anesthetized and 0.03 mg/kg of buprenorphine and 1 mL of saline were administered subcutaneously. Mice were then transferred to a stereotaxic apparatus (Harvard Apparatus) and maintained at 2-2.5% isoflurane. The mice were placed on a heating pad and a probe was rectally inserted to maintain a body temperature of 37°C. Prior to laser exposure, mice were intraperitoneally injected with 10 mg/kg of photoactive Rose Bengal (Sigma) followed by a 5-minute delay to allow the dye to enter circulation. Skin at the top of the head was surgically cut to expose the skull and then the sensorimotor cortex was targeted using stereotaxic coordinates (3 cm above, mediolateral + 0.24 mm from Bregma). The skull of the mice was exposed to a laser (Beta Electronics, wavelength: 532 nm) for 15 minutes. For recovery of post-operative pain, buprenorphine was administered to all animals prior to ischemic damage.

### Behavioral Testing

#### Bederson Scale and Neurological Scoring Scale

Following stroke, animals subsequently exhibit a variety of neurological deficits. The Bederson scale is a global neurological assessment that was developed to measure neurological impairments following stroke (Bederson et al., 1986). Tests include forelimb flexion, resistance to lateral push and circling behavior. A severity scale of 0-3 is used to assess behavioral deficits after stroke. This scoring scale is a simple way to reveal basic neurological deficits. Ischemic animals will have significantly more neurological deficits than non-ischemic animals, resulting in a higher score (Bederson et al., 1986).

#### Accelerating Rotarod

A standard accelerating rotarod apparatus (Harvard Apparatus) was used to measure walking movements and balance previously described (Balkaya et al., 2013; Jadavji et al., 2017, 2015). Thirty centimeters above the ground, mice were placed on a rotating rod 3 cm in diameter and 6 cm wide in which the speed gradually increases from 4 to 60 RPM over 8 min. When mice fall off the rotarod, a digital sensor records the latency in seconds. An average of three trials per mouse was taken with an inter trial interval of 5 minutes.

#### Forepaw Placement

To measure spontaneous forepaw usage, mice were placed in a 19 cm high, 14 cm diameter cylinder, and the placement of their forepaws on the cylinder wall during natural exploratory rearing behaviors was recorded using a digital camera for frame-by-frame analysis (Jadavji et al., 2017; Theoret et al., 2016). During a rear, the first forepaw placement on the wall was recorded as impaired, non-impaired, or both.

#### Ladder Beam

The ladder rung apparatus was composed of two Plexiglas walls. Each wall contained holes located at the bottom edge of the wall; the holes could be filled with metal bars. The entire apparatus was placed atop two standard mouse housing cages. The performance was video-recorded from the side, with the camera positioned at a slight ventral angle so that all 4 limbs could be recorded at the same time (Farr et al., 2006).

All video recordings were analyzed frame-by-frame. Each step was scored according to the quality of limb placement as previously described (Farr et al., 2006). For analysis of foot placement accuracy, the number of errors in each session was counted. The error score was calculated from the total number of errors and the number of steps for each limb (Farr et al., 2006).

### Total Homocysteine and Choline Metabolite Measurements

At the time of euthanization, blood was collected by cardiac puncture in EDTA coated tubes, centrifuged at 7000g for 7 min at 4°C to obtain plasma. Liver tissue was also removed at the same time and samples were stored at −80°C, until time of analysis. Total homocysteine (tHcy), *S*-adenosylmethionine, S-adenosylhomocysteine, methionine, cystathionine, betaine, and choline were measured in plasma and liver by liquid chromatography tandem mass spectrometry (LC-MS/MS) as previously described (Ducros et al., 2006).

### Brain Tissue Processing

Some mice were perfused and fixed brain tissue was sectioned using a cryostat at 30 μm and slide mounted in serial order. There were six slides full of brain tissue sections of the damaged area per mouse and each animal had a minimum of four sections that were used for quantification. ImageJ (NIH) was used to quantify ischemic damage volume by measuring the area of damaged tissue (Jensen, 2013).

### Immunofluorescence Experiments

Brain tissue was used in immunofluorescence analysis to assess apoptosis, using active caspase-3 (1:100, Cell Signaling Technologies). All brain sections were stained with a marker for neuronal nuclei, NeuN (1:200, AbCam). Primary antibodies were diluted in 0.5% Triton X and incubated with brain tissue overnight at 4°C. The next day, brain sections were incubated in Alexa Fluor 488 or 555 (Cell Signaling Technologies) and secondary antibodies were then incubated at room temperature for 2 hours and stained with 4’, 6-diamidino-2-phenylindole (DAPI) (1:1000, Thermo Fisher Scientific). The stains were analyzed using a microscope (Zeiss) and all images were collected at the magnification of 400X.

In brain tissue within the ischemic region, co-localization of active caspase-3 with NeuN labeled neurons were counted and averaged per animal. A positive cell was indicated by co-localization of the antibodies of interest located within a defined cell. Cells were distinguished from debris by identifying a clear cell shape and intact nuclei (indicated by DAPI and NeuN) under microscope. All cell counts were conducted by two individuals blinded to treatment groups. The number of positive cells were counted in three brain sections per animal. For each section, three fields were analyzed. The number of positive cells were averaged for each animal.

### Metabolomics

Some mice were euthanized using CO_2_ overdose. Cecum and brain tissue (damaged and non-damaged) was microdissected and immediately frozen for analysis. Prior to LC-MS/MS targeted measurement, frozen tissue supernatant samples were first thawed overnight under 4°C. Afterward, 20 mg of each thawed sample were placed in a 2 mL Eppendorf vial. The initial step for protein precipitation and metabolite extraction was performed by adding 500 μL MeOH and 50 μL internal standard solution (containing 1,810.5 μM ^13^C_3_-lactate and 142 μM ^13^C_5_-glutamic acid). The mixture was then vortexed for 10 s and stored at −20°C for 30 min, followed by centrifugation at 14,000 RPM for 10 min at 4°C. The supernatants (450 μL) were collected into new Eppendorf vials and dried using a CentriVap Concentrator. The dried samples were reconstituted in 150 μL of 40% PBS/60% ACN and centrifuged again at 14,000 RPM at 4°C for 10 min. Afterward, 100 μL of supernatant was collected from each sample into an LC autosampler vial for subsequent analysis. A pooled sample, which was a mixture of all experimental samples, was used as the quality control (QC) sample and injected once every 10 experimental samples.

The targeted LC-MS/MS method used here was modeled after that developed and used in a growing number of studies (Bapat et al., 2021; Jasbi et al., 2021, 2019a, 2019b; Wang et al., 2019). Briefly, all LC-MS/MS experiments were performed on an Agilent 1290 UPLC-6490 QQQ-MS system. Each supernatant sample was injected twice, 10 µL for analysis using negative ionization mode and 4 µL for analysis using positive ionization mode. Both chromatographic separations were performed in hydrophilic interaction chromatography mode on a Waters XBridge BEH Amide column (150 × 2.1 mm, 2.5 µm particle size, Waters Corporation, Milford, MA). The flow rate was 0.3 mL/min, auto-sampler temperature was kept at 4°C, and the column compartment was set to 40°C. The mobile phase was composed of Solvents A (10 mM NH_4_OAc, 10 mM NH_4_OH in 95% H_2_O/5% ACN) and B (10 mM NH_4_OAc, 10 mM NH_4_OH in 95% ACN/5% H_2_O). After an initial 1 min isocratic elution of 90% B, the percentage of Solvent B decreased to 40% at t = 11 min. The composition of Solvent B was maintained at 40% for 4 min (t = 15 min), after which the percentage of B gradually went back to 90%, to prepare for the next injection. The mass spectrometer was equipped with an electrospray ionization (ESI) source. Targeted data acquisition was performed in multiple-reaction-monitoring (MRM) mode. The whole LC-MS system was controlled by Agilent MassHunter Workstation software. The extracted MRM peaks were integrated using Agilent MassHunter Quantitative Data Analysis software.

### Data Analysis and Statistics

Behavioral and brain tissue data were analyzed by two individuals that were blinded to experimental treatment groups. GraphPad Prism 6.0 was used to analyze behavioral testing, plasma tHcy measurements, lesion volume, immunofluorescence staining, and choline measurements. In GraphPad Prism 6.0, two-way ANOVA analysis was performed when comparing the mean measurement of both sex and dietary group for behavioral testing, one-carbon metabolites, lesion volume, and immunofluorescence staining. Significant main effects of two-way ANOVAs were followed up with Tukey’s post-hoc test to adjust for multiple comparisons. MetaboAnalyst 5.0 was used to analyze brain and cecum metabolomics data (Pang et al., 2021). Measured metabolites were mapped to the Kyoto Encyclopedia of Genes and Genomes (KEGG) reference database for *Mus musculus* to assess pathway enrichment. Metabolomics data were normalized to tissue weight and log-transformed prior to both univariate and multivariate comparisons. All data are presented as mean ± standard error of the mean (SEM). Statistical tests were performed using a significance level of 0.05.

## Results

### One-Carbon Metabolites

Homocysteine levels were measured in plasma and liver tissues 5 weeks after ischemic stroke. Females maintained on a vit. B12 def. diet had higher levels of plasma (*p* = 0.008) and liver (*p* = 0.0012) total homocysteine levels compared to controls (Table 1).

**Table 1.**
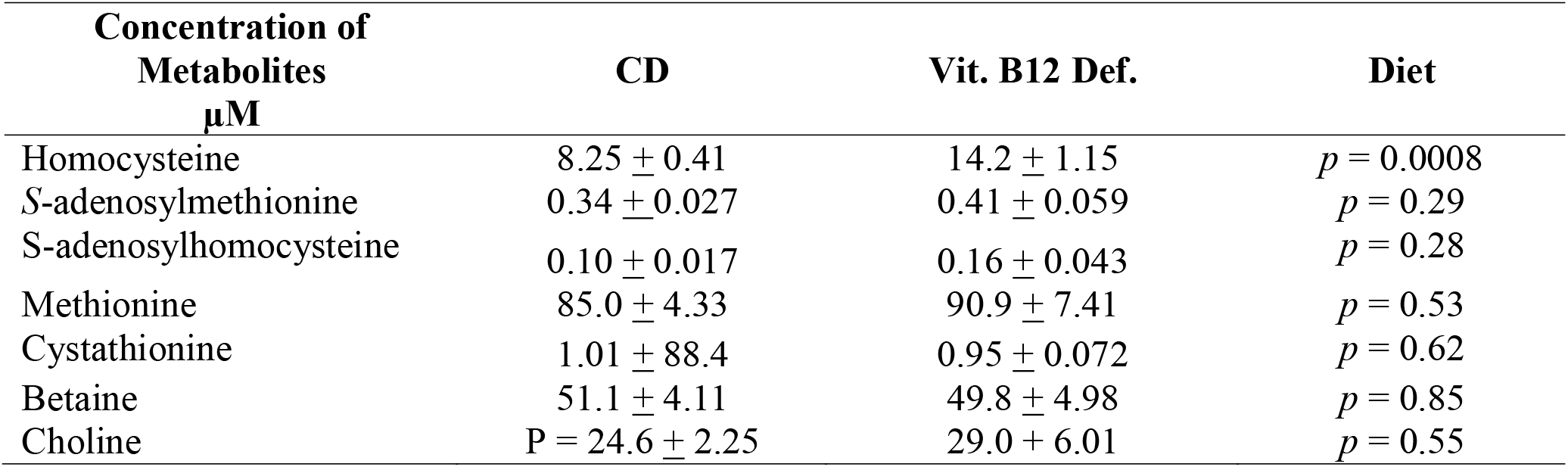
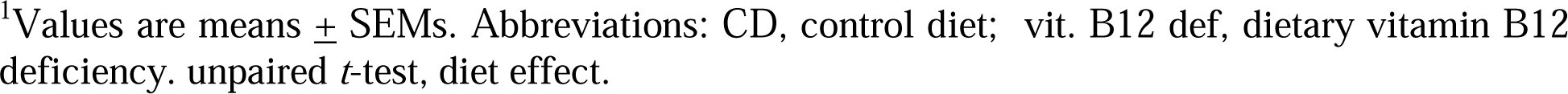
One-carbon metabolite concentrations in plasma of control and vit. B12 def. diet females.^1^

| Concentration of<br>Metabolites<br>$\mu\text{M}$ | CD | Vit. B12 Def. | Diet |
| --- | --- | --- | --- |
| Homocysteine | $8.25 \pm 0.41$ | $14.2 \pm 1.15$ | $p = 0.0008$ |
| S-adenosylmethionine | $0.34 \pm 0.027$ | $0.41 \pm 0.059$ | $p = 0.29$ |
| S-adenosylhomocysteine | $0.10 \pm 0.017$ | $0.16 \pm 0.043$ | $p = 0.28$ |
| Methionine | $85.0 \pm 4.33$ | $90.9 \pm 7.41$ | $p = 0.53$ |
| Cystathionine | $1.01 \pm 88.4$ | $0.95 \pm 0.072$ | $p = 0.62$ |
| Betaine | $51.1 \pm 4.11$ | $49.8 \pm 4.98$ | $p = 0.85$ |
| Choline | $P = 24.6 \pm 2.25$ | $29.0 \pm 6.01$ | $p = 0.55$ |

<sup>1</sup>Values are means ± SEMs. Abbreviations: CD, control diet; vit. B12 def, dietary vitamin B12 deficiency. unpaired *t*-test, diet effect.

There were no differences in other one-carbon metabolites in liver tissue except higher levels of choline (*p* = 0.02) in vitamin B12 deficient animals compared to control diet animals (Table 2).

**Table 2.** One-carbon metabolite concentrations in liver tissue of control and vit. B12 def. diet females.^1^

| Concentration of<br>Metabolites<br>$\mu\text{mol/g}$ | CD | Vit. B12 Def. | Diet |
| --- | --- | --- | --- |
| Homocysteine | $6.39 \pm 0.34$ | $8.98 \pm 0.40$ | $p = 0.001$ |
| S-adenosylmethionine | $547 \pm 94.1$ | $797 \pm 213$ | $p = 0.37$ |
| S-adenosylhomocysteine | $425 \pm 21.7$ | $454 \pm 37.1$ | $p = 0.53$ |
| Methionine | $5033 \pm 442$ | $4746 \pm 362$ | $p = 0.62$ |
| Cystathionine | $256 \pm 20.5$ | $249 \pm 24.9$ | $p = 0.85$ |
| Betaine | $123 \pm 11.2$ | $117 \pm 12.8$ | $p = 0.73$ |
| Choline | $257 \pm 35.2$ | $373 \pm 21.6$ | $p = 0.02$ |

<sup>1</sup>Values are means ± SEMs. Abbreviations: CD, control diet; vit. B12 def, dietary vitamin B12 deficiency.

### Ischemic Damage Volume

Ischemic damage volume was measured in brain tissue 5 weeks following photothrombosis damage. Representative images of cresyl violet stained brain tissue are shown in Figure 1A from control and vitamin B12 deficient animals. Quantification of ischemic damage shows that vit. B12 def. animals had greater damage volume compared to controls (Figure 1B; *p* = 0.01).

**Figure 1.**
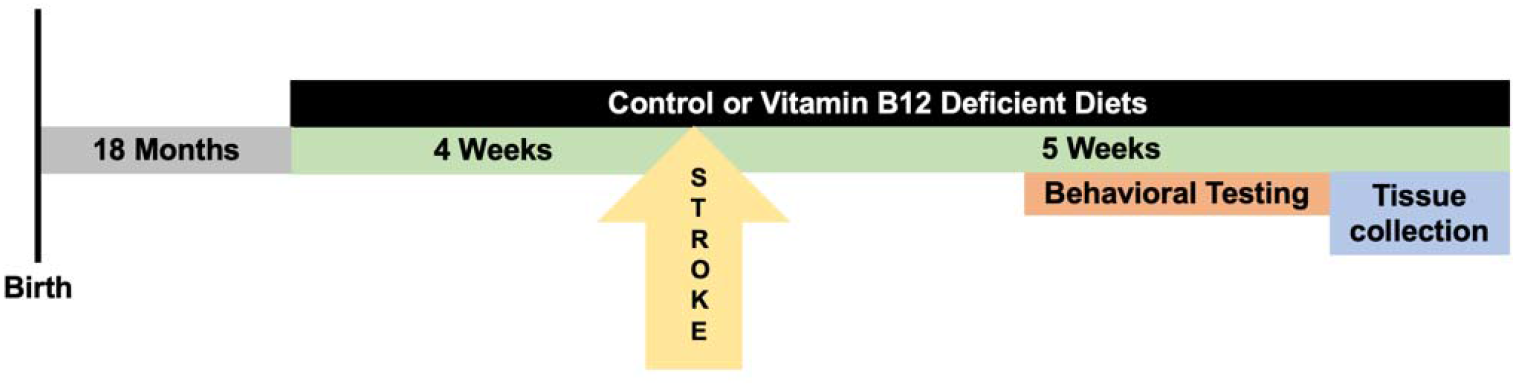
Timeline of experimental manipulation. Female C57Bl/6J mice arrived from Jackson Laboratories 10-months-old, and after one week of acclimation, animals were placed on a control or vitamin B12 deficient diet for four weeks. Following the four weeks, ischemic stroke was induced using the photothrombosis model in the sensorimotor cortex. After stroke animals were maintained on respective diets for 4 additional weeks, motor function of the mice was measured using the accelerating rotarod and forepaw placement tasks. At the completion of *in vivo* experiments animals were euthanized, and brain and liver tissue, as well as plasma was collected for further analysis.

### Neuronal Apoptosis

Within the ischemic damage region we measured levels of neuronal apoptosis using active caspase-3 and NeuN. Representative images of immunofluorescence staining are shown in Figure 2A from control and vit. B12 def. diet animals. Quantification of apoptosis shows that vitamin B12 deficient animals had more apoptosis compared to controls (Figure 2B, *p* = 0.05).

**Figure 2.**
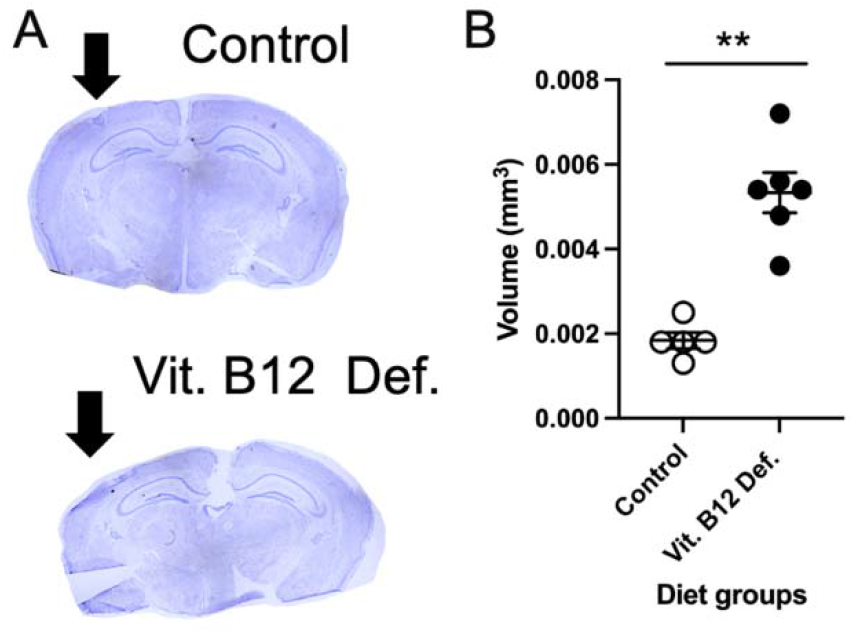
Impact of dietary vitamin B12 deficiency and ischemic stroke on damage volume. Representative cresyl violet image (A) and ischemic damage volume quantification (B). Depicted are means of ± SEM of 4 to 5 mice per group.

### Metabolomic Measurements

#### Brain Mitochondria

In total, 34 compounds were reliably detected in samples (i.e., QC coefficient of variation (CV) < 20%). A two-factor heatmap showing normalized relative abundance of captured metabolites by group and lesion status is visualized in Supplementary Figure S1. Pearson’s correlation and clustering analysis revealed mainly positive correlations, with few negative correlations identified (Supplementary Figure S2). Specifically, strong positive correlations (*r* ≥ 0.5) were found within metabolites of phenylalanine metabolism, branched-chain amino acid (BCAA) metabolism, and metabolites of the TCA cycle. Meanwhile, strong negative correlations (*r* ≤ −0.5) were observed between amino acids and TCA cycle metabolites. An orthogonal partial least squares-discriminant analysis (OPLS-DA) was performed using the total set of 34 detected metabolites to assess brain mitochondria metabolite profiles (Supplementary Figure S3A) and permutation testing with 100 iterations was performed to assess model fit (Supplementary Figure S3B). The OPLS-DA scores plot showed notable separation between control and vitamin B12 deficient groups and the model showed good predictive capacity (*Q*^*2*^ = 0.435) and high explanatory capacity (*R*^*2*^ = 0.815). Importantly, permutation testing showed the model was not overfit to the data (perm. *p* < 0.05). A two-way MANOVA with all measured metabolites was performed to test main effects of ischemic and non-ischemic stroke, as well as B12 deficiency. Although no significant main effect by stroke type was observed (i.e., ischemic vs. non-ischemic stroke), a significant main effect of vit. B12 def. was observed in eight captured metabolites: phenylalanine (*p* = 0.012), malate (*p* = 0.013), tyrosine (*p* = 0.014), aspartate (*p* = 0.019), alanine (*p* = 0.027), valine (*p* = 0.029), isoleucine (*p* = 0.039), and fumarate (*p* = 0.047). Normalized box plots of significant metabolites are provided in Figure 3A. Unsupervised principal component analysis (PCA) was performed with the subset of eight significant metabolites and showed appreciable separation between control and vitamin B12 deficient animals, with more than 83% of total variance explained (Supplementary Figure S4). Furthermore, receiver operating characteristic (ROC) analysis was performed with the eight significant metabolites; we noted two significant metabolites (phenylalanine and tyrosine) had univariate area under curve (AUC) estimates > 0.90 (Supplementary Figure S5), suggesting these two metabolites to be highly accurate indicators of B12 deficiency. Phenylalanine showed an AUC = 0.988, (95% CI: 0.9-1.0, sensitivity = 0.9, specificity = 1.0), while tyrosine showed a similarly high AUC = 0.925 (95% CI: 0.75-1.0, sensitivity = 0.8, specificity = 0.9).

**Figure 3.**
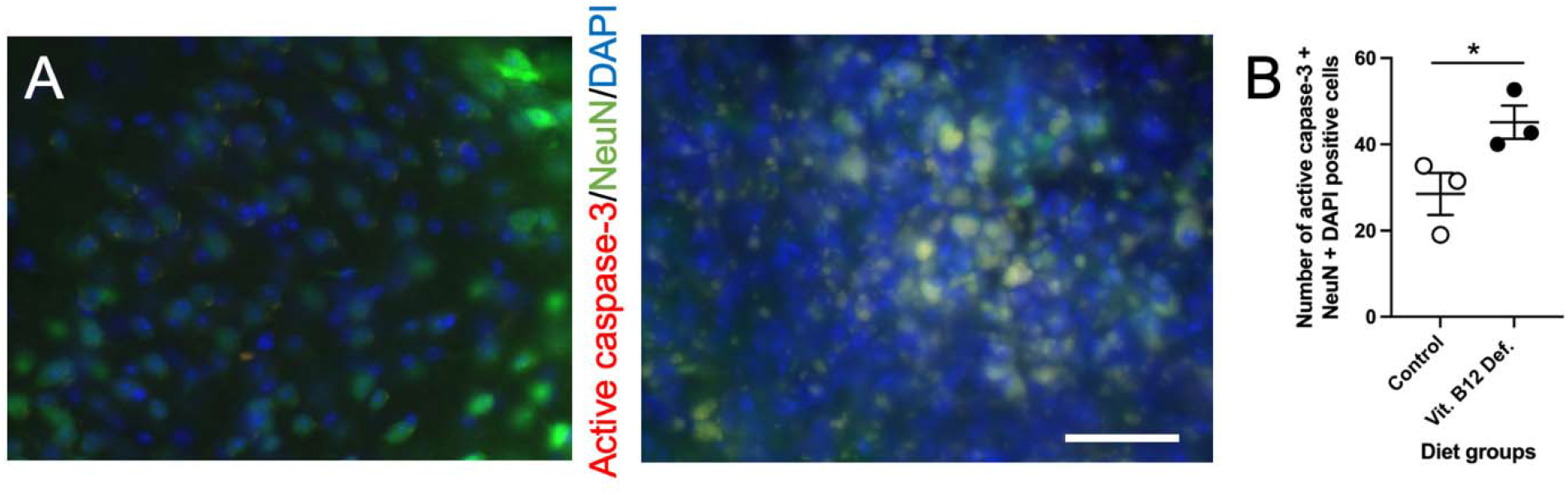
Impact of dietary vitamin B12 deficiency and ischemic stroke on neuronal active caspase-3 cell counts. Representative images of immunofluorescence staining with positive semi-quantitative spatial colocalization of active caspase-3 with neuronal nuclei (NeuN) and 4′,6-diamidino-2-phenylindole (DAPI) (A). Quantification of active capsapse-3, NeuN, and DAPI cell counts (B). Depicted are means of ± SEM of 3 mice per group. The scale bar = 50 μm.

Additionally, measured metabolites were annotated using the *Mus musculus* KEGG reference database and pathway enrichment analysis was performed between B12 deficient and control animals using a global test of relative distance betweenness centrality. Results are visualized as a scatter plot in Figure 3B, showing significance by pathway impact. One pathway was shown to be significantly impacted between groups: phenylalanine, tyrosine, and tryptophan biosynthesis (*p* = 0.007). Importantly, all metabolites in this pathway were measured by the metabolomic assay, and two showed significance (impact = 1.0).

Results of our pathway analysis showed significant alterations (FDR *q* < 0.05) in nine canonical murine pathways (Figure 4B): 1) alanine, aspartate and glutamate metabolism, 2) glycine, serine, and threonine metabolism, 3) cysteine and methionine metabolism, 4) valine, leucine, and isoleucine biosynthesis, 5) lysine biosynthesis, 6) arginine and proline metabolism, 7) histidine metabolism, 8) phenylalanine, tyrosine and tryptophan biosynthesis, and 9) tryptophan metabolism. Notably, these nine pathways are constitutive of the aminoacyl tRNA biosynthesis superpathway. As such, we have summarized our findings across subpathways of greatest impact with regard to aminoacyl tRNA biosynthesis in Figure 4. As can be seen, metabolomic analysis of brain mitochondria implicates dysregulated aminoacyl tRNA biosynthesis as a primary driver of impaired motor function following stroke, particularly in constituent pathways containing phenylalanine, tyrosine, aspartate, alanine, fumarate, isoleucine, and valine. Importantly, these effects are observed at both the metabolite (0.01 ≤ *p* ≤ 0.05) and pathway (0.01 ≤ *q* ≤ 0.02) levels.

**Figure 4.**
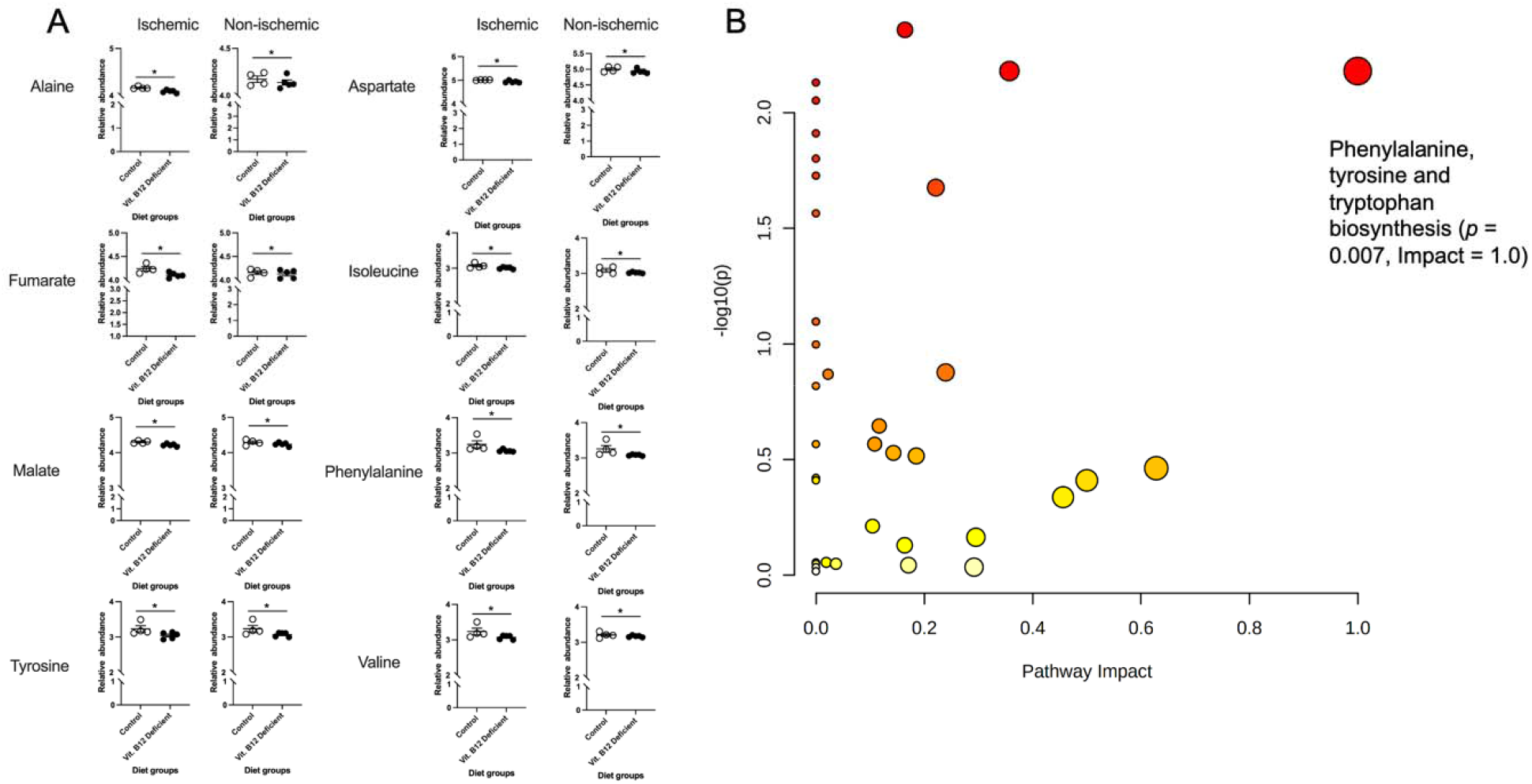
Impact of dietary vitamin B12 deficiency and ischemic stroke on mitochondria metabolomics in ischemic and non-ischemic brain tissue. Significance testing and enrichment analysis of metabolite data. Data were normalized to sample weight and log-transformed prior to analysis. (A) Significant between diet group metabolites (regardless of ischemic damage) as determined by two-way MANOVA: alanine (*p* = 0.027), aspartate (*p* = 0.019), fumarate (*p* = 0.047), isoleucine (*p* = 0.039), malate (*p* = 0.013), phenylalanine (*p* = 0.012), tyrosine (*p* = 0.014), valine (*p* = 0.030),. (B) Pathway enrichment analysis performed using KEGG canonical pathways. Enriched pathways with *p* < 0.05 and impact > 0.50 have been labeled.

#### Cecum

In cecum samples, 158 metabolites were reliably detected after filtering (i.e., QC CV < 20%). Independent samples *t*-testing of metabolites between vitamin B12 deficient and control diet groups revealed two significant metabolites: creatinine (*p* = 0.04) and methylmalonic acid (*p* = 0.04). Normalized box plots of these significant metabolites are provided in Figure 5A, and are further visualized as a heatmap between groups in Figure 5B. Fold change (FC) analysis (B12 deficient/control diet) was also performed to assess the magnitude of changes in metabolites. Although non-significant, three metabolites (3-aminobutyric acid, shikimic acid, and 4-hydroxybenzoic acid) showed FC > 2 and three metabolites (DOPA, 3-phenyllactic acid, and DOPA) showed FC < 0.5. These results are summarized and displayed in Figure 5C.

**Figure 5.**
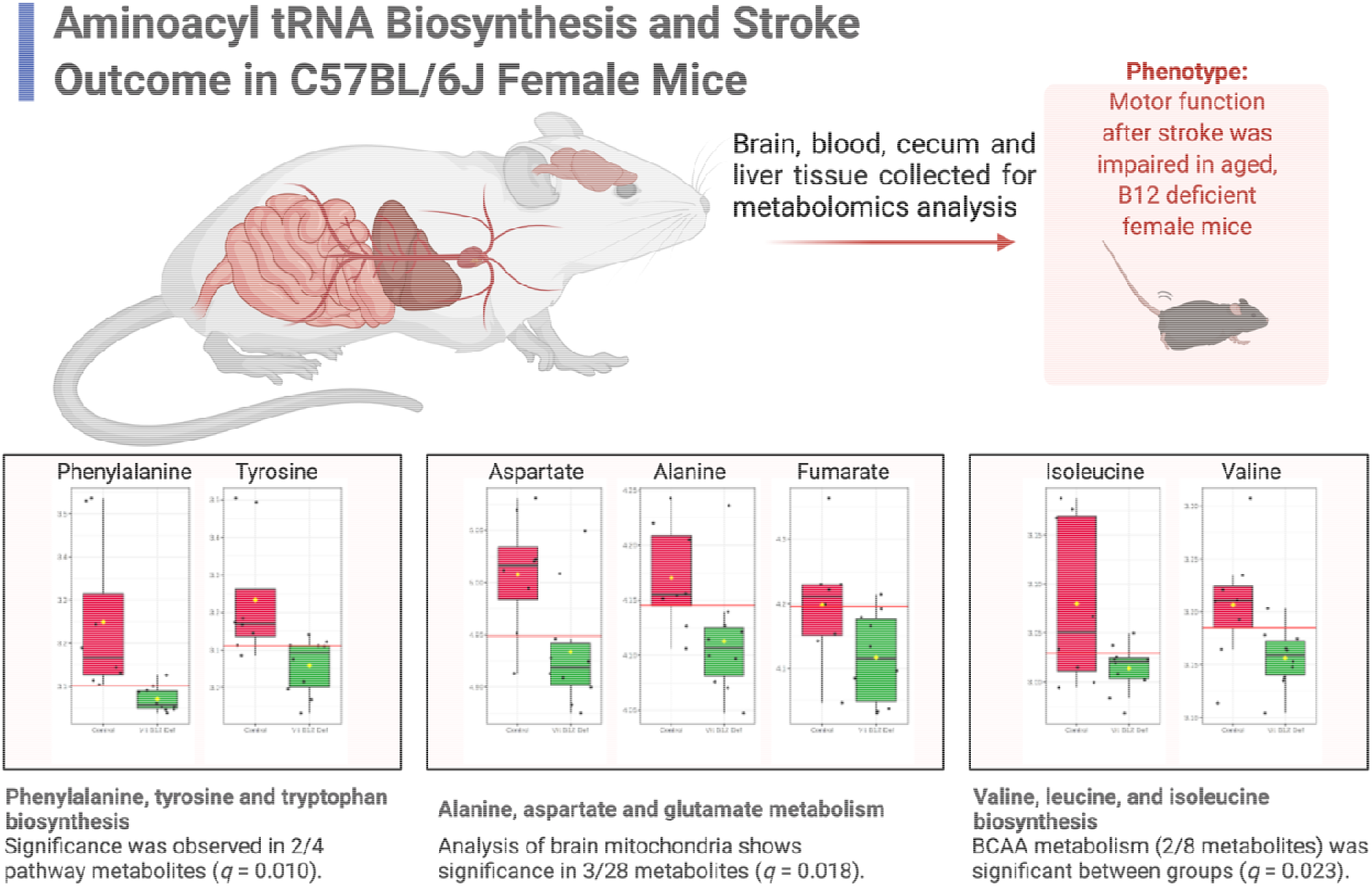
Pathway analysis results summarized in the context of aminoacyl tRNA biosynthesis. Metabolomic analysis of brain mitochondria suggests involvement of phenylalanine, tyrosine, and tryptophan biosynthesis, alanine, aspartate and glutamate metabolism, and valine, leucine, and isoleucine biosynthesis in impaired motor function following stroke (all metabolites *p* < 0.05). Predicted functional significance (*q*) adjusted for FDR. BCAA, branched-chain amino acids.

### Behavioral Testing

#### Neuro deficit Score

Four weeks after damage, we measured neuro deficit in animals. Vit. B12 def animals had a higher severity score on the vertical screen test compared to control diet animals (Figure 6A, *p* = 0.02). There was no difference in the postural reflex (*p* = 0.50) and forelimb placing (*p* = 0.60) tests.

**Figure 6.**
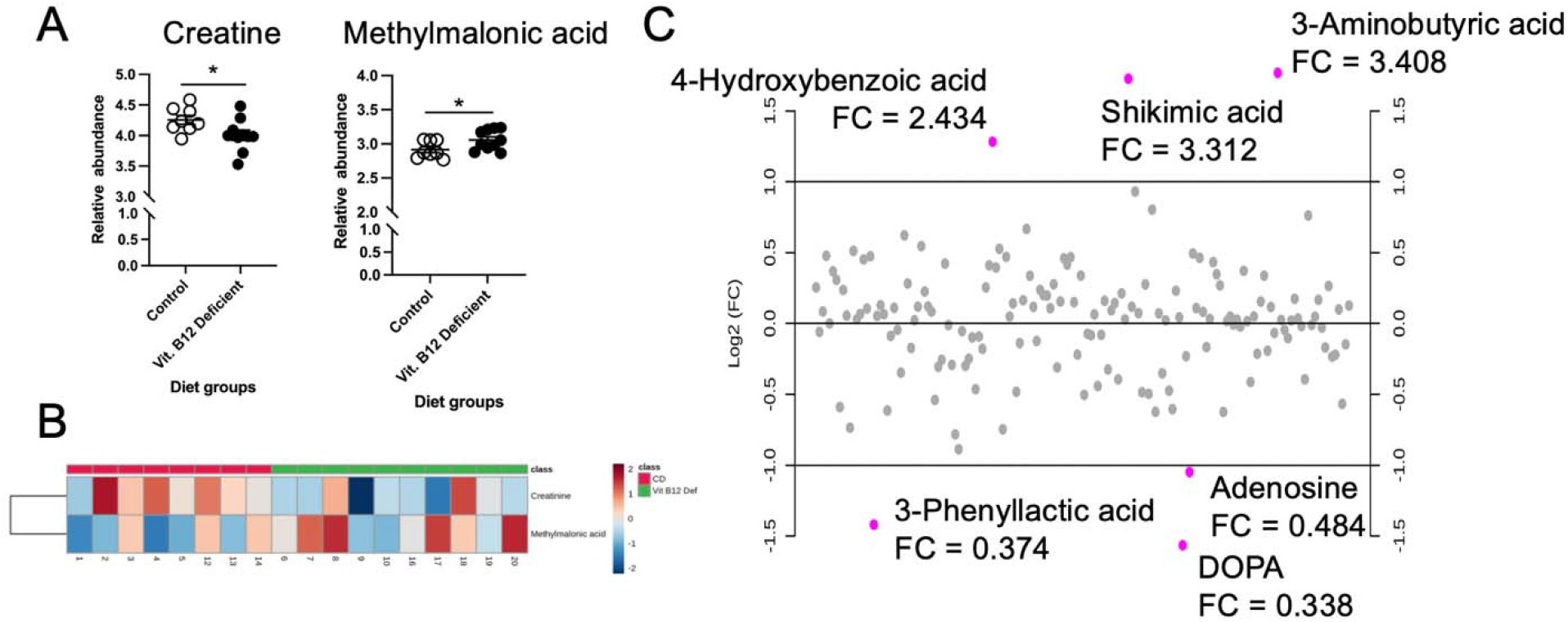
Impact of dietary vitamin B12 deficiency and ischemic stroke on cecum metabolomics. Significance testing, heatpmap visualization, and fold change (FC) analysis of metabolite data. Data were normalized to sample weight and log-transformed prior to analysis; samples were unpaired, equal group variances assumed, and parametric test performed. (A) Significant between-group metabolites as determined by independent samples *t*-test: creatinine (*p* = 0.04), methylmalonic acid (*p* = 0.04). (B) Heatmap display of creatinine and methylmalonic acid. (C) FC analysis (performed as B12 deficient/control diet) showed three metabolites with FC > 2 (4-hydroxybenzoic acid, shikimic acid, 3-aminobutyric acid) and three metabolites with FC < 0.5 (3-phenyllactic acid, DOPA, adenosine).

**Figure 7.**
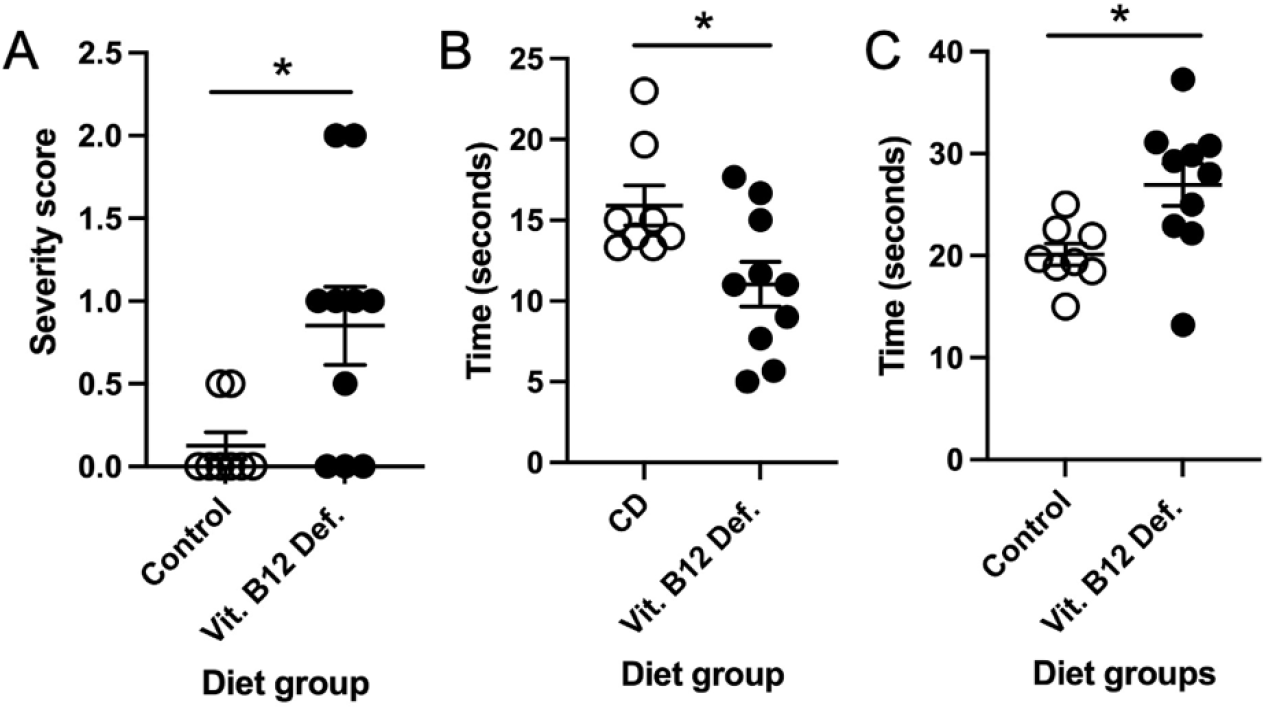
Impact of vitamin B12 deficiency on motor function after ischemic stroke. Neurodeficit score, vertical screen test severity score (A). Latency to fall off the accelerating rotarod (B). Amount of time to cross the ladder beam (C). Eight to ten mice per group. * p < 0.05, un-paired t-test.

#### Accelerating Rotarod

After ischemic stroke, vit. B12 def. female mice were not able to stay on the accelerating rotarod as long as control animals (Figure 6B, *p* = 0.021). There was no difference in the revolutions per minute between groups (*p* = 0.90).

#### Forepaw Placement

We measured whether there was a difference between forepaw usage after stroke. No difference between vit. B12. def. and controls were observed in impaired (*p* = 0.22) and non-impaired (*p* = 0.098) forepaw usage.

#### Ladder Beam

There was no difference in the movement score of the impaired (*p* = 0.72) and non-impaired (*p* = 0.66) limbs. The number of errors made while crossing the ladder did not differ between control and vit. B12. def. animals for impaired (*p* = 0.17) or non-impaired (*p* = 0.42) limbs. The vit. B12. def. diet animals did take longer to cross the ladder beam compared to control animals (Figure 6C, *p* = 0.02).

## Discussion

A vitamin B12 deficiency results in increased risk of stroke and worse stroke outcome in patients (Yahn et al., 2021), although the mechanisms through which this occurs remain unknown. In this study we investigated the mechanisms through which a vitamin B12 deficiency impacts stroke outcome in aged female mice. After ischemic stroke, vitamin B12 deficient animals had increased levels of total homocysteine in plasma and liver. There were no changes in one-carbon metabolites in plasma, but in liver tissue choline levels were increased in vitamin B12 deficient animals. Vitamin B12 deficient females had larger ischemic damage volume in brain tissue and more apoptosis within the ischemic region. Our study is the first to describe mitochondria metabolite changes in brain tissue of control and vitamin B12 deficient aged female animals after ischemic stroke. In the cecum we describe changes in methylmalonic acid and creatinine; pathway enrichment analysis showed dysfunction in B12 transport. Analysis of mitochondrial metabolomics in brain tissue showed significant decreases in phenylalanine, malate, tyrosine, aspartate, alanine, valine, isoleucine, and fumarate in response to a vitamin B12 deficiency and not ischemic damage. Meanwhile, pathway analysis showed significant, widespread dysfunction in phenylalanine, tyrosine, and tryptophan biosynthesis. Stroke functional outcome was also measured, and showed that vitamin B12 deficient females performed worse on the vertical screen test and had impaired coordination and balance.

We confirmed the dietary vitamin B12 deficiency in animals by measuring levels of plasma and liver homocysteine. Our metabolite analysis also revealed that levels of methylmalonic acid were increased in the cecum. Clincally, both increased levels of homocysteine and methylmalonic acid are markers for a vitamin B12 deficiency (Langan and Goodbred, 2017). Others have used the same vitamin B12 deficient diet used in this study and have demonstrated that it results in a deficiency (Bernard et al., 2018), and our data builds upon this literature.

Using aged mice (19-months-old), our preclinical data confirms clinical observations that a vitamin B12 deficiency results in worse ischemic stroke outcome in aged females (Ahmed et al., 2019; Yahn et al., 2021). In our study we targeted the sensorimotor cortex using the photothrombosis model, which is a model of ischemic stroke. The model of stroke has been reported to result in reproducible damage (Carmichael, 2006, 2005; Fluri et al., 2015; Lee et al., 2004). We have also observed the reproducible damage in the present study as well as others (Abato et al., 2020; Jadavji et al., 2018, 2019, 2017). The mice used in the present study are defined as aged animals modeling aspects of older humans (Fluri et al., 2015), making this a valid preclinical model. We report reduced levels of creatine in the cecum of vitamin B12 deficient mice, which has been reported to delay neurodegeneration (Smith et al., 2014). Reduced levels of creatine in the cecum may be playing a role in the increased neurodegeneration we observe in the brain tissue of vitamin B12 deficient animals after ischemic stroke.

The data from this study provide insight into how the brain tissue of vitamin B12 deficient aged females respond to ischemic stroke. We have shown more damage in the brain tissue after ischemic stroke and increased levels of neuronal apoptosis within the damaged region. Our metabolite measurements in healthy and ischemic brain tissue revealed that metabolites of the tricarboxylic acid (TCA) cycle (malate and fumarate) are reduced in vitamin B12 deficient animals. The TCA cycle has been implicated in the response to hypoxia (Martínez-Reyes and Chandel, 2020). The induction of stress like ischemic stroke may push more neurons to die faster.

Vitamin B12 plays a number of roles within the cell, it is involved in the methylation of homocysteine and generation of *S*-adenosylmethionine (SAM), a global methyl donor. Additionally, in the mitochondria, vitamin B12 is involved in the regulation of mitochondrial metabolism. Recent data suggests that vitamin B12 plays an important antioxidant role in the body and, when levels are reduced, there is more oxidation present (An et al., 2021; van de Lagemaat et al., 2019). The increased levels of oxidation lead to the activation of nuclear factor-erythroid factor 2-related factor (Nrf-2) (An et al., 2021), an important transcription factor involved in cellular response to oxidative stress. Ischemic stroke also results in oxidative stress and increased levels of Nrf-2 (Farina et al., 2021). In our previous study, we showed that supplementation with B vitamins, including vitamin B12, results in increased levels of Nrf-2 and superoxide dismutase 2 (SOD2) (Jadavji et al., 2017). Another study has described that vitamin B12 can act as a scavenger for reactive oxygen specific (ROS) after renal ischemia and reperfusion, as well as reduce inflammation (“The beneficial role of vitamin B12 in injury induced by ischemia-reperfusion,” 2021).

In conclusion, our stroke outcome data corroborates the current clinical data, specifically, that a vitamin B12 deficiency results in worse stroke outcome. The driving force of this reduced outcome maybe changes in mitochondrial metabolism and aminoacyl tRNA biosynthesis leading to more neurodegeneration in brain tissue after ischemic stroke.

## Supporting information

DOI: 10.17632/mwkvc9kw7p.1.

## Abbreviations

CD: Control diet
CVD: Cardiovascular disease
1C: one carbon
vit. B12 def.: vitamin B12 deficiency

## Acknowledgments

None

## Supplementary Information

Supplementary materials are available online.

**Figure S1** Two-factor heatmap of 34 brain mitochondria metabolites by group/lesion.

**Figure S2** Correlation and clustering analysis of brain mitochondria metabolites.

**Figure S3** OPLS-DA and permutation testing with 34 brain mitochondria metabolites.

**Figure S4** PCA scores plot between study groups using eight significant metabolites.

**Figure S5** Univariate ROC analysis of phenylalanine and tyrosine for B12 deficiency.

## Data Availability Statement

The data that supports the findings of this study have been deposited to Mendeley data and are publicly available at DOI: 10.17632/mwkvc9kw7p.1. and are available in the supplementary material of this article.

