## Supplementary material for "Ischemic stroke and dietary vitamin B12 deficiency in old-aged females impaired motor function, increased ischemic damage size, and changed metabolite profiles in brain and cecum tissue": DOI: 10.17632/mwkvc9kw7p.1.

---

### Reduced stroke outcome in old-aged female mice maintained on a dietary vitamin B12 deficiency

**Page S-3, Figure S1** Two-factor heatmap showing normalized relative abundance of 34 captured metabolites of brain mitochondria by group (control vs. vitamin B12 deficient) and lesion status (lesion vs. non-lesion).

**Page S-4, Figure S2** Pearson's correlation and clustering heatmap between 34 detected study metabolites from bran mitochondria. Correlation and significance cutoffs were set to $r \geq |0.5|$ and $p < 0.05$, respectively.

**Page S-5, Figure S3** Orthogonal partial least squares-discriminanat analysis (OPLS-DA) performed with 34 captured metabolites of the brain mitochondria. (A) OPLS-DA scores plot between control and vitamin B12 deficient groups ($Q^2$ = 0.435, $R^2$ = 0.815). (B) Permutation testing with 100 iterations (perm. $p < 0.05$).

**Page S-6, Figure S4** Unsupervised principal component analysis (PCA) performed with the subset of eight significant metabolites of the brain mitochondria. PC1 and PC2 explain more than 83% of total variance.

**Page S-7, Figure S5** Receiver operating characteristic (ROC) analysis of vitamin B12 deficiency in brain mitochondria. (A) Univariate area under curve (AUC) of phenylalanine (AUC = 0.988, 95% CI: 0.9-1.0, sensitivity = 0.9, specificity = 1.0). (B) Standard box plot of normalized phenylalanine measurements showing optimal cutoff (red line) and group means (yellow diamonds). (C) Univariate AUC of tyrosine (AUC = 0.925, 95% CI: 0.75-1.0, sensitivity = 0.8, specificity = 0.9). (D) Standard box plot of normalized tyrosine measurements showing optimal cutoff (red line) and group means (yellow diamonds).

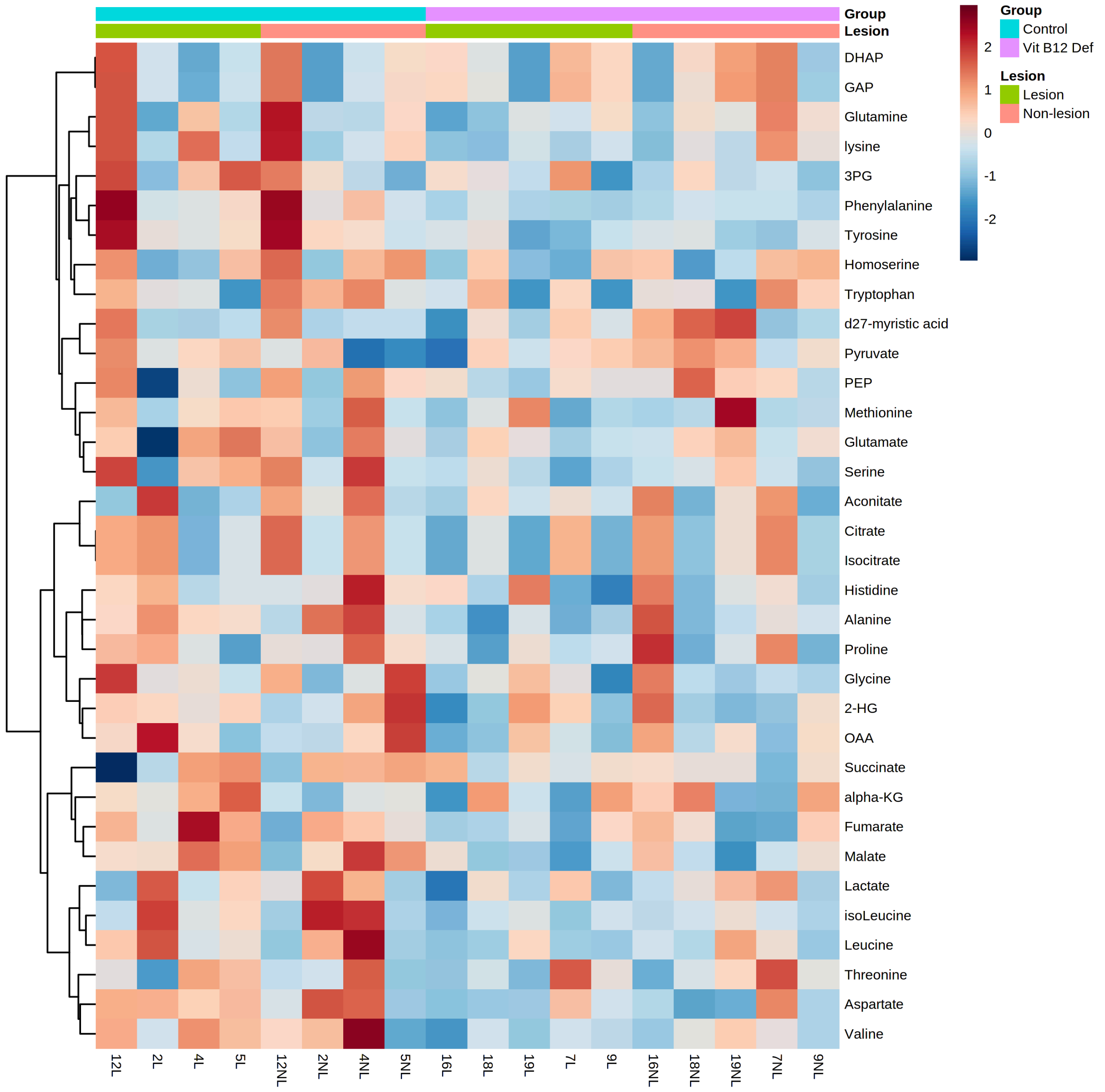


**Figure S1** Two-factor heatmap showing normalized relative abundance of 34 captured metabolites of brain mitochondria by group (control vs. vitamin B12 deficient) and lesion status (lesion vs. non-lesion).

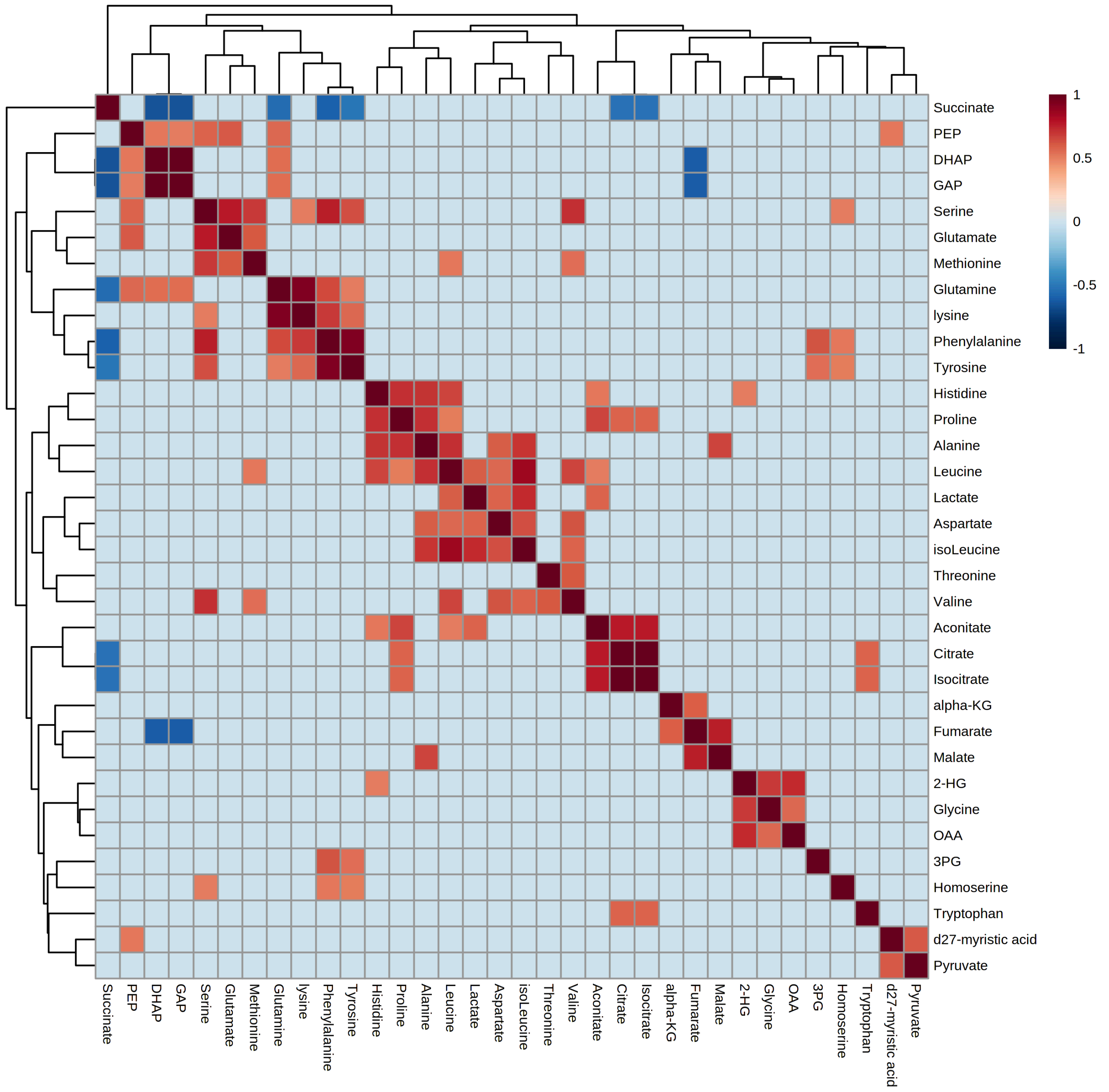


**Figure S2** Pearson's correlation and clustering heatmap between 34 detected study metabolites from bran mitochondria. Correlation and significance cutoffs were set to $r \geq |0.5|$ and $p < 0.05$, respectively.

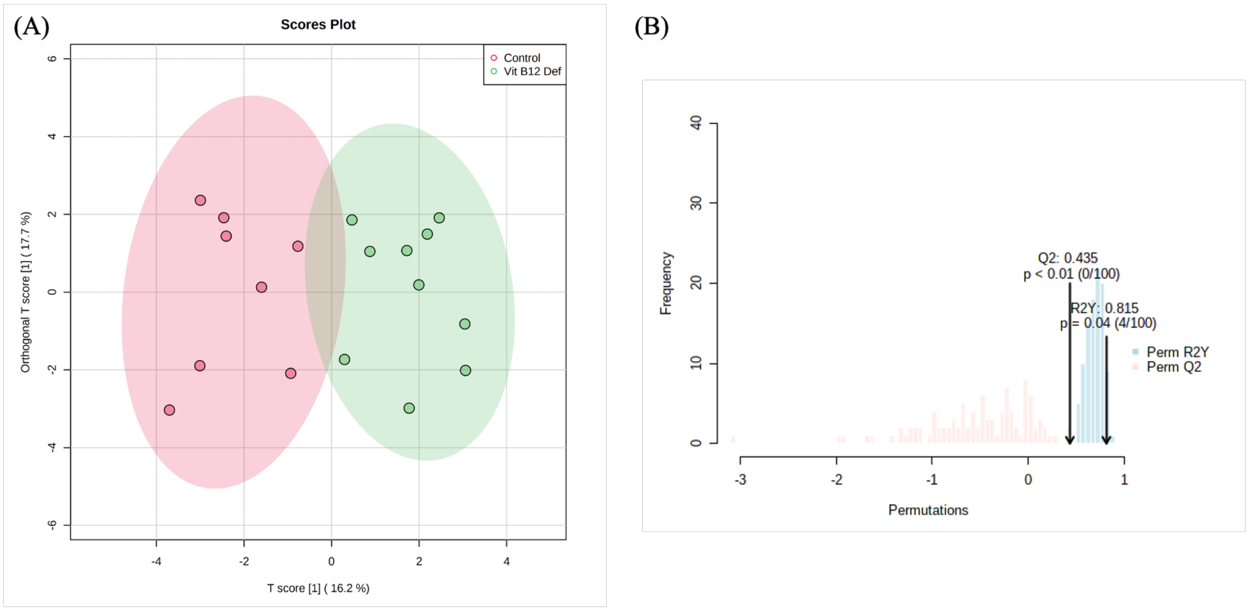


**Figure S3** Orthogonal partial least squares-discriminanat analysis (OPLS-DA) performed with 34 captured metabolites of the brain mitochondria. (A) OPLS-DA scores plot between control and vitamin B12 deficient groups ($Q^2 = 0.435$, $R^2 = 0.815$). (B) Permutation testing with 100 iterations (perm. $p < 0.05$).

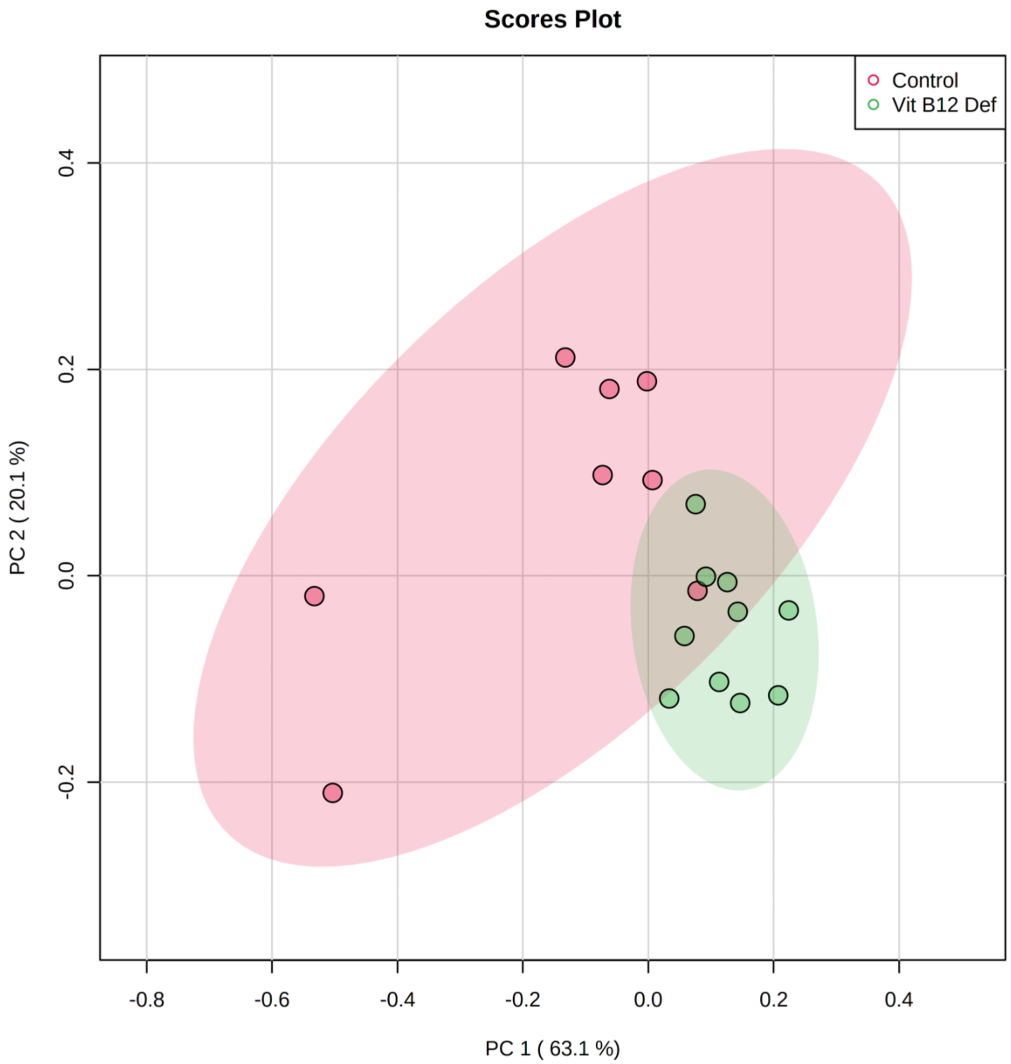


**Figure S4** Unsupervised principal component analysis (PCA) performed with the subset of eight significant metabolites of the brain mitochondria. PC1 and PC2 explain more than 83% of total variance.

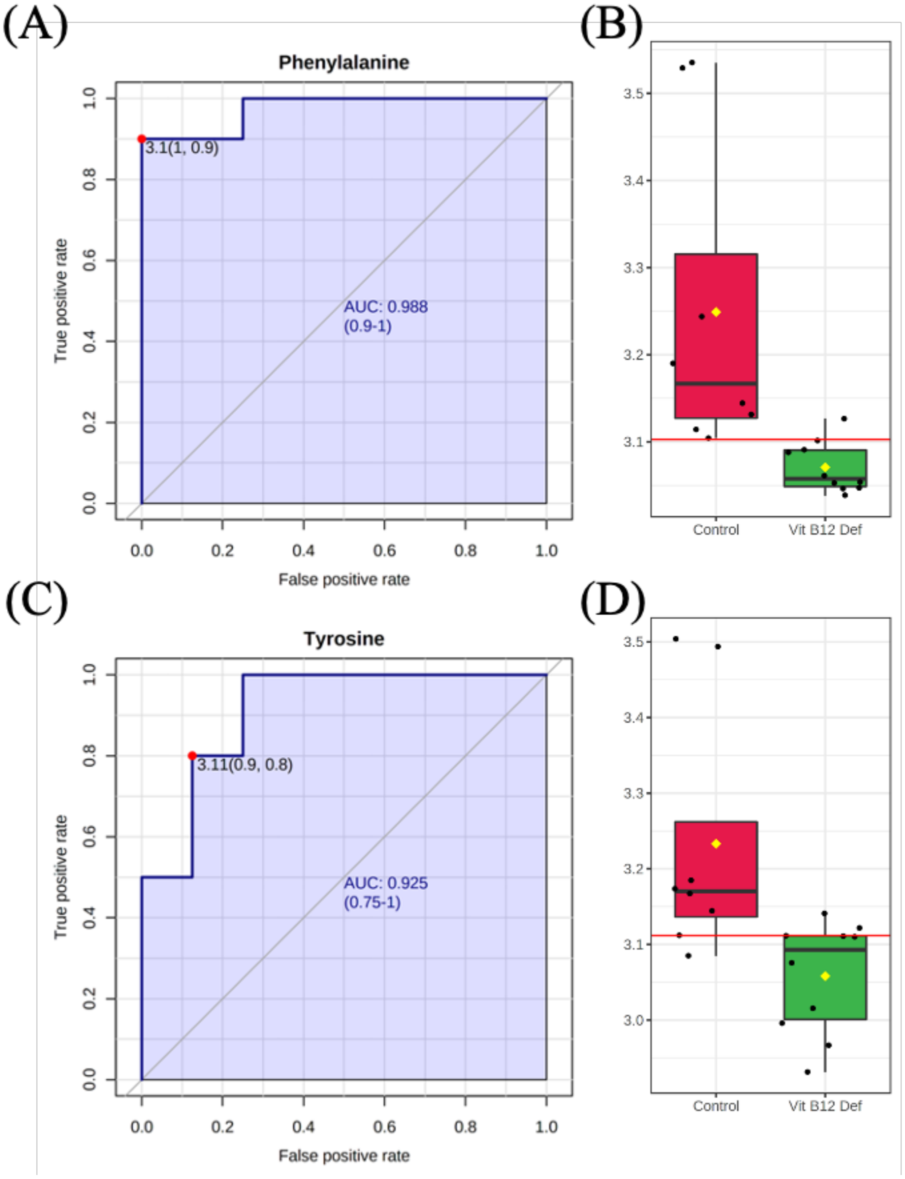


**Figure S5** Receiver operating characteristic (ROC) analysis of vitamin B12 deficiency in brain mitochondria. (A) Univariate area under curve (AUC) of phenylalanine (AUC = 0.988, 95% CI: 0.9-1.0, sensitivity = 0.9, specificity = 1.0). (B) Standard box plot of normalized phenylalanine measurements showing optimal cutoff (red line) and group means (yellow diamonds). (C) Univariate AUC of tyrosine (AUC = 0.925, 95% CI: 0.75-1.0, sensitivity = 0.8, specificity = 0.9). (D) Standard box plot of normalized tyrosine measurements showing optimal cutoff (red line) and group means (yellow diamonds).
